# A Minimally Invasiveness Hybrid Brain-Computer Interface: A Distributed, Scalable and Evolvable Architecture for Whole Brain Access

**DOI:** 10.64898/2026.07.14.738604

**Authors:** Zhe Li, Na Liu, Li Wan, Mengting Liu, Chenguang Wu

## Abstract

Brain-computer interfaces face a fundamental trade-off between the signal fidelity and stimulation precision of noninvasive systems and the surgical burden and scalability of invasive systems. Non-invasive BCIs suffer from low signal quality and poor stimulation accuracy due to the skull barrier and the variability introduced by the scalp and skull. Existing invasive BCIs rely on traumatic surgical procedures or brain-penetrating electrodes, which limits their spatial extensibility, application, and patient acceptance. Here, we introduce a minimally invasive hybrid BCI architecture that uses the skull as a distributed interface layer rather than treating it solely as a barrier. The hybrid BCI comprises four integrated components: (1) the safe and smart micro-hole craniotomy; (2) distributed microelectrodes subcutaneously implanted in micro-holes in the skull with the distal end in contact with the dura; (3) an external bi-directional wearable headset for coupling, recording, stimulation, and channel selection; and (4) an AI-assisted planning and control agent. Animal studies have shown that micro-holes with a diameter of 300-800 μm can be safely and conveniently prepared at any predefined locations across the skull without impairing the dura. In vivo experiments on rats demonstrate that the hybrid BCI with skull-implanted microelectrodes evidently increases resting-state spectral power and improves the signal-to-noise ratio of somatosensory and steady-state visual evoked responses compared to the scalp EEG; the computational modelling shows that distributed skull-dura microelectrodes can increase the intracranial electrical field strength and steer focused temporal-interference fields towards predefined deep brain targets. These findings will lay a solid foundation for future endeavors in wireless integration, safety evaluation and clinical benefits of the hybrid BCI. In summary, we propose the hybrid BCI as a distinct minimally invasive BCI paradigm with the great potential as a distributed, scalable, and upgradable neural interface that can expand the clinical application of minimally invasive BCI techniques.

## 1. Introduction

Brain-computer interfaces (BCIs) establish a direct functional link between neural activity and external devices. They can record neural signals (“read”), deliver electrical, optical, or acoustic energy (“write”), or combine these operations in a closed loop, with applications in communication, motor and sensory restoration, neurorehabilitation, and neuromodulation. The central translational challenge is not simply to maximize signal quality or minimize invasiveness in isolation, but to achieve a clinically acceptable balance among information fidelity, procedural burden, chronic stability, spatial coverage, and system upgradability.

Clinical BCI technologies span a continuum of invasiveness. In March 2026, China’s National Medical Products Administration approved NEO, an epidural implantable BCI for upper-limb motor rehabilitation, marking an important regulatory milestone for semi-invasive BCI translation. Other approaches include subdural cortical arrays, endovascular electrodes, and intracortical microelectrode arrays. These systems can provide high-fidelity neural access than the scalp EEG, but their broader use is constrained by the surgical burden, chronic interface stability, device explantation or revision requirements, and the difficulty of extending interfaces across multiple brain regions.

Experience with other implanted neurotechnologies illustrates that technical eligibility does not guarantee clinical uptake. In deep brain stimulation (DBS), only a minority of patients with Parkinson’s disease (1.6% to 4.5%) meet the eligibility criteria for DBS^1^; and many eligible patients (up to 60-70%) decline surgery^2,3^, due to the fear of surgery^4^, uncertain benefits^5^, or concerns of recovery^5^. Similarly, cochlear implantation remains underused among eligible adults. The global acceptance of cochlear implant is only about 10%^6,7^; even in high-income countries, its acceptance rate is still below 20%^8^. Although these technologies are not equivalent to BCI, they highlight how procedural burden, risk-benefit uncertainty, and patient preference can limit adoption of implanted neural devices. A scalable BCI platform must therefore address both neural performance and long-term usability.

Non-invasive BCIs use scalp-mounted wearable sensors and offer ease of deployment. However, the skull attenuates, filters, and spatially smooths electrical potentials measured at the scalp^9–11^. These effects arise from the skull’s relatively low^12^ and frequency-dependent conductivity^13^, heterogeneous microstructure^14^, and variable thickness^15^. For instance, the electrical conductivity of skull is only ∼20 mS/m; in comparison, the electrical conductivity for scalp is 410 mS/m^12^. Motion, muscle activity, electrode-skin impedance, and placement variability further reduce inter-session consistency. Advanced signal-processing and artificial intelligence methods can improve denoising and neural decoding; however, they cannot fully recover spatial and physiological information that is not captured at the sensor level.

The skull also constraints precise transcranial neuromodulation^16^. In transcranial electrical stimulation, due to the “shunt effect”, a substantial fraction of the applied current is shunted through the scalp and skull, reducing intracranial field strength, and limiting the focality. It was reported that only ∼10% of stimulation current applied on the scalp actually entered the brain^17^, making it difficult to stimulate specific brain regions via non-invasive strategies. Temporal-interference stimulation offers a potential route to stimulate deep brain targets, but targeting remains sensitive to tissue heterogeneity, electrode placement, and safety-limited current amplitudes^18^. Ultrasound and optical neuromodulation use different propagation mechanisms, yet they also require accurate modeling and control of energy transmission through heterogeneous cranial tissues^19,20^.

These limitations motivate an interface that approaches the dura without penetrating the dura or the brain tissue and that keeps the progressively evolving electronic system outside the body. A skull-based micro-interface could reduce the impedance and spatial uncertainty introduced by the skull while avoiding conventional craniotomy and intracranial leads. However, this concept requires a reproducible method for creating micro-channels through the hard skull without impairing the underlying dura, a stable and biocompatible electrode-dura interface, and a reliable external coupling architecture.

Existing mechanical and laser-based approaches can create small cranial openings, but endpoint control, thermal injury, and selective dura protection remain important engineering challenges. Owing to the high-strength skull^21,22^ and the complex brain-dura interface^23,24^, it is hard to achieve safe micro-hole craniotomy with existing material removal techniques such as mechanical drilling^25,26^ or laser drilling^27,28^. Use of a micro-drill bit would result in a high probability of tool fracture; a rotating micro-drill bit would also damage dura. As for micro-hole craniotomy by laser machining, presence of blood or tissue fluid will disturb the optical coherence tomography (OCT) signals for real-time depth control, making the cutting through detection method unreliable. In the in vivo test performed by researchers from Medical University of Vienna^29^, dura or even cortex damage was hard to avoid in robot-driven laser beam craniotomy, owing to the uncertainty in identifying the end point of craniotomy.

Here, we propose the minimally invasive hybrid BCI architecture based on four integrated elements, including (1) the acupuncture-like safe and smart micro-hole craniotomy, (2) distributed microelectrodes implanted in micro-holes at predefined positions in the skull, with the distal electrode in contact with the dura and the proximal end beneath the scalp, (3) an external bidirectional wearable headpiece for signal acquisition, stimulation, channel selection, and power or data coupling; and (4) an AI-assisted planning and control agent for electrode implantation planning, signal decoding, electric-field optimization, and future closed-loop control of the hybrid BCI. In this context, the “hybrid” refers to the functional coupling between the chronically implanted distributed microelectrodes with the external wearable and upgradable electronics.

The present study evaluates three enabling technical layers of this architecture. First, we characterize the mechanism, thermal control, and endpoint detection of the ultrasonic micro-hole craniotomy. Second, we test whether skull-implanted microelectrodes improve resting and evoked neural signal recordings in comparison with the conventional scalp EEG. Third, we use computational modeling to examine whether distributed skull-dura microelectrodes can increase intracranial field strength and focality, and steer temporal-interference fields toward cortical and deep brain targets. The study does not yet demonstrate a fully integrated wireless, chronic, or closed-loop human system; rather, it establishes fundamental cornerstones of the hybrid BCI by verifying the feasibility of the micro-intervention access, high-fidelity recording, and field-shaping principles. Conceptually, the architecture occupies a distinct design space between scalp-based noninvasive BCIs and intracranial invasive BCIs and will enable distributed, incrementally expandable neural interface for whole brain access.

## 2. Results

### The ultra-minimally invasive micro-hole craniotomy technique and device

We have developed a technique for the safe and smart micro-hole craniotomy (S^2^MHC) without material removal, for the implementation of minimally invasive hybrid BCI via micro-intervention. This technique was proposed by Dr. Zhe Li around 2015 for the first time. In this study, it has been pushed to a mature state for clinical application. Exploiting the unique structural (e.g., the porous architecture) and material (e.g., organic and inorganic compositions) properties of the skull, this technique utilizes high-amplitude ultrasonic vibration to locally deform bio-composite’s microporous architecture, achieving safe, smart, high-quality and low-force (< 500 mN) micro-hole craniotomy (diameter: 300-800 μm) on the high-strength skull with minimal thermal effect (maximum temperature < 42 ℃; postoperative healing in 2 weeks), in a convenient and minimally invasive manner similar to subcutaneous injection on the skin. Micro-tunnels generated at pre-defined locations across the skull would function as micro-I/O ports for communicating with the brain. Implanting microelectrodes into the brain and using a wearable headset coupled with these micro-implants, a hybrid BCI could be established for acquiring high-quality neural signal or precise neural stimulation (Fig. 1a).

**Figure 1.**
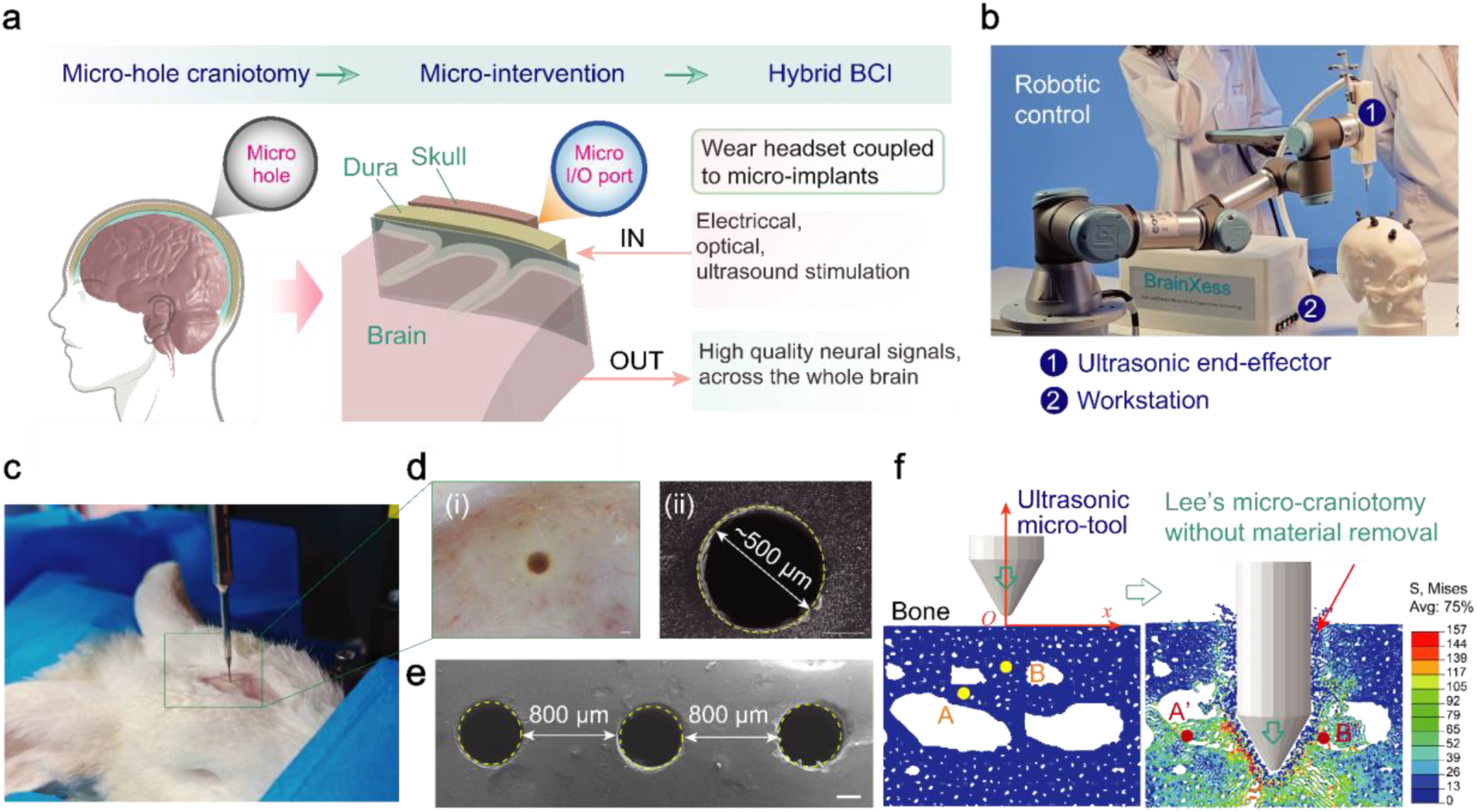
The micro-hole craniotomy technique and device. (a) The technical pipe-line of micro-hole craniotomy, micro-intervention, and the minimally invasive hybrid BCI. Through micro-I/O ports prepared on the skull, micro-intervention can be conveniently performed by implanting microelectrode into these micro-holes, generating a minimally invasive passage for the hybrid BCI. (b) The micro-hole craniotomy device, consisting of an ultrasonic end-effector and a workstation; an interchangeable micro-tool is mounted on the end-effector for micro-hole craniotomy without material removal. (c) Micro-hole craniotomy performed at an anesthetized New Zealand white rabbit. The ultrasonic vibration assisted micro-hole craniotomy method, or Lee’s micro-hole craniotomy, first proposed by Dr. Zhe Li around 2015. (d) The optical and SEM mages showing the high-quality micro-hole generated in the skull in (c). (e) Scanning electron microscopic (SEM) image showing micro-holes with a diameter of 500 μm diameter can be densely created in the skull at an interval less than 1 mm. Scale bar in d-e: 200 μm. (f) Simulation of the ultrasonic vibration assisted micro-hole forming process without material removal using the smooth particle hydrodynamics method; A and B are two representative bone tissue particles, while A’ and B’ are their corresponding position in the micro-hole craniotomy process.

For this purpose, we have developed a smart micro-hole craniotomy device, consisting of an ultrasonic end-effector and a workstation (Fig. 1b). The end-effector can be mounted onto a robotic arm for micro-hole craniotomy across the whole skull along a pre-planned trajectory. Multifaceted challenges in areas including device development, micro-hole craniotomy mechanism and performance, thermal necrosis management, and termination detection at the irregular skull-dura interface are effectively addressed through a holistic approach. With these endeavors, high-quality micro-holes could be precisely generated on the skull. Fig. 1c-e demonstrates the generation of an array of micro-holes (diameter: 500 μm) at an interval of 800 μm through a minimally invasive process without material removal (see the simulation results in Fig. 1f).

### Acupuncture-like micro-hole craniotomy with minimal thermal effects

The mechanism of ultrasonic vibration assisted micro-hole craniotomy was investigated with simulation (using the smooth particle hydrodynamics method^30,31^, Fig. 2a) and finite-depth craniotomy tests on freshly harvested skull bone from rabbits (by advancing the ultrasonic micro-tool to a prescribed depth *h* and then withdraw; Fig. 2b). Upon tool-bone contact, the ultrasonic micro-tool would microscopically damage bone tissue (see bone tissue debris in the “crater” in Fig. 2b at a feed depth of *h* = 100 μm). Along with tool feeding, high frequency hammering of the ultrasonic tool would repeatedly “forge” fractured bone debris (see the smooth inner surface at *h* = 200 μm in Fig. 2b); skull bone’s micro-porous architecture would be plastically deformed and squeezed sidewards around the conical tip, creating a high-quality micro-hole (good circularity and smooth inner surface) in the high-strength skull bone without material removal (*h* = 400 μm in Fig. 2b). In the micro-hole craniotomy process, a layer of densely compacted bone tissue was plastically formed around the micro-hole.

**Figure 2.**
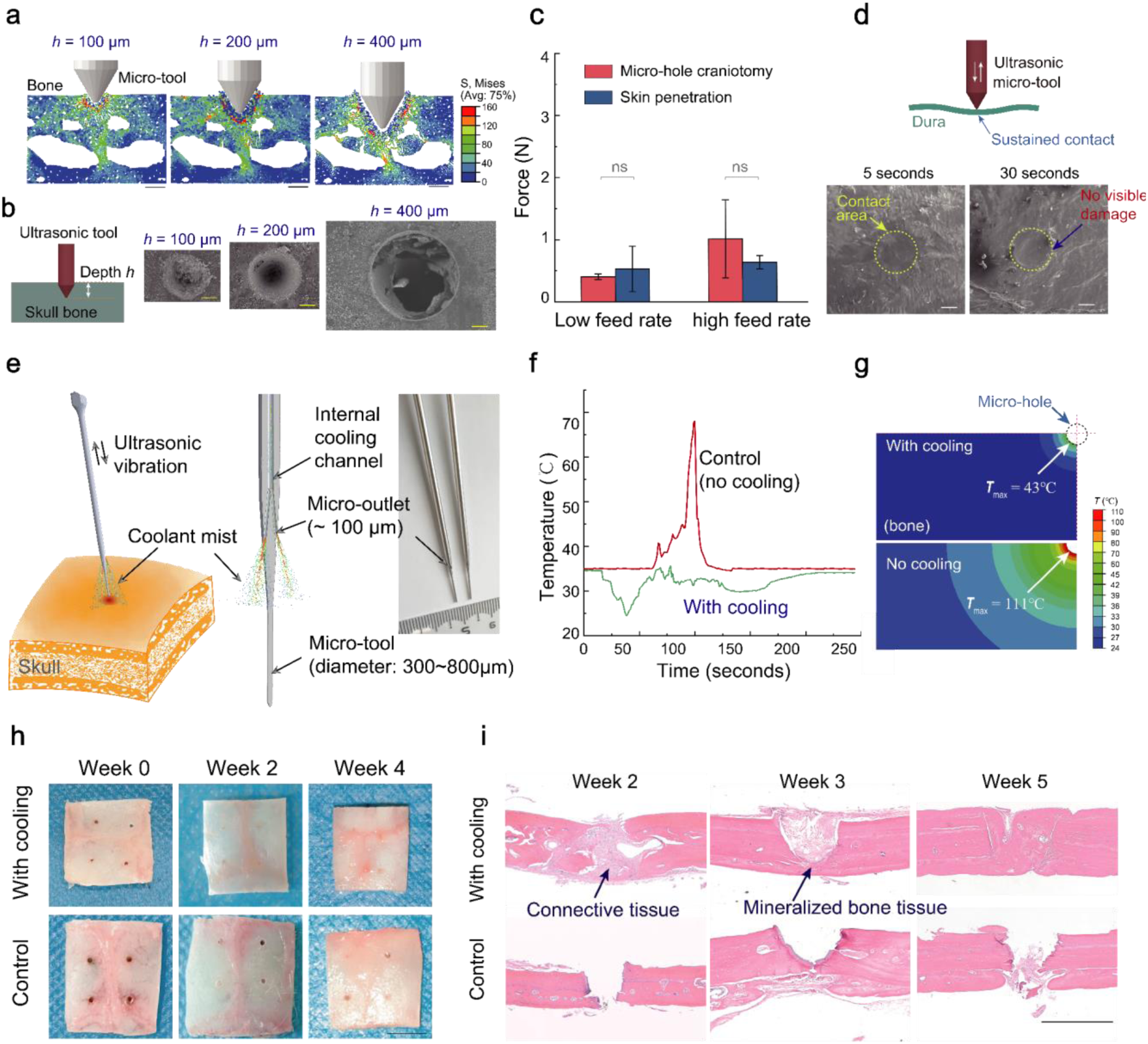
Acupuncture-like high quality micro-hole craniotomy with a low force and minimal thermal effects. (a) Simulation of the micro-hole craniotomy at different feed depth *h*; scale bar: 200 μm. (b) SEM images of the finite-depth micro-hole craniotomy test; the ultrasonic micro-tool was fed to a given depth *h* in skull and withdrawn; scale bar: 100 μm. (c) Low-force micro-hole craniotomy. The micro-hole craniotomy force is comparable to skin penetration performed on rat skin with a subcutaneous injection needle (diameter: ∼ 500 μm; *n* = 4); ns: no significant difference (*P*<0.01). (d) Safe micro-hole craniotomy at the skull-dura interface. SEM images show that after sustained contact with the ultrasonic micro-tool, the dura is intact without visible damage; scale bar: 100 μm. Rabbit bone and dura samples were used in c-d. (e) Micro-hole craniotomy with effective cooling; micro-channels could be machined in the long and slender micro-tool with micro-outlets prepared on the side-wall for cooling; without this design, low-thermal effects would be difficult to achieve. (f) Representative temperature-time curves measured at 0.5 mm off the hole wall with optimized cooling and without cooling. (g) Simulated temperature contours around the micro-hole with cooling and without cooling. In f-g *in vitro* tests were performed on freshly harvested rabbit skull bone half-immersed in the water-bath to a physiological temperature of 37℃. (h-i) Skull samples harvested from rats at different time after micro-hole craniotomy and the corresponding histological images after hematoxylin-eosin (H&E) staining; scale bar in h: 4.0 mm; scale bar in **i**: 300 μm.

Performance of the micro-hole craniotomy technique was further evaluated. The micro-hole craniotomy force can be as low as ∼ 500 mN, which is even smaller than the force for subcutaneous injection on soft tissue (Fig. 2c), despite the significant difference in elastic modulus between skull and skin (skull bone’s Young’s modulus is 4 orders of magnitude higher than skin^32^). In the control group without ultrasonic vibration, the micro-tool could be forced into skull bone, but resulting in a significantly large force, deteriorated hole morphology and unpredictable tool fracture, which is disastrous and unacceptable in clinical applications.

The ultrasonic micro-hole craniotomy technique is extremely safe at the skull-dura interface when the skull bone is totally penetrated. The device can selectively deform the hard bone tissue while effectively protecting the soft tissue with negligible machining capability (such as the dura beneath skull). As shown in Fig. 2d, sustained tool-dura contact by the ultrasonic on did not cut or penetrate the dura mater. Even after 30 seconds of continuous contact, no visible tear or damage could be observed on the dura under SEM (Fig. 2d). This selective machining capability can ensure safe micro-hole craniotomy at the skull-dura interface.

The ultrasonic micro-hole craniotomy technique has minimal thermal effects on the skull bone (Fig. 2e-g). In the micro-hole craniotomy process without cooling, heat generation generated by the friction between the plastically deformed bone tissue and the ultrasonic micro-tool would cause the temperature in the bone tissue around the micro-hole to rise to about 110 ℃ (Fig. 2g). If left untreated, this would lead to thermal necrosis, impairing post-operation recovery. To address this problem, we have developed an effective cooling method. With advanced micro-machining techniques, a co-axial micro-channel for internal cooling was prepared in the long slender tool, with micro-outlets (size: about 100 μm) prepared on the sidewall at the distal end. Pressurized saline coolant delivered through the micro-outlet could be atomized by the ultrasonic vibration and applied to the craniotomy site (Fig. 2e). With optimized cooling, the bone temperature measured at 0.5 mm off the hole wall was only ∼34.2℃ (Fig. 2f), effectively preventing heat build-up and keeping the bone tissue around the micro-hole at a low temperature. Incorporating the experimentally measured temperature data into the thermal conduction model, the maximal temperature at hole wall was predicted to be about 43℃, significantly lower than that without cooling (Fig. 2g). Without cooling, irreversible thermal necrosis would be induced when bone tissue was exposed to a temperature of >47°C for one minute^33^.

Micro-hole craniotomy with low thermal effects was confirmed *in vivo*. At different weeks after micro-hole craniotomy on SD rats, bone samples were harvested for histological staining^34^(Fig. 2h-i). Due to excessive heat generation during micro-hole craniotomy without cooling, the micro-hole was not healed even after 5 weeks (Fig. 2h). In contrast, with optimized cooling, micro-holes were able to heal in 2 weeks and the hole cavity was filled by regenerated soft tissue; bone mineralization was even observed at Week 3 (Fig. 2i). Thus, we show that with optimized cooling, the micro-hole craniotomy could allow rapid postoperative recovery (e.g., in 2 weeks). Micro-hole craniotomy and rapid recovery significantly minimize the invasiveness of the micro-intervention operation.

### Smart control to enhance the safety of micro-intervention at the skull-dura interface

Precise detection of the craniotomy termination at the skull-dura interface is crucial for safe micro-hole craniotomy (Fig. 3a). For this purpose, we have developed a smart control system consisting of a hardware unit and a breakthrough detection algorithm. The breakthrough detection algorithm was developed with multi-mode information (Fig. 3a), which were filtered and processed for control purposes^35^. The ultrasonic device could be further integrated with a robotic arm and an optical navigation system (Fig. 3b), allowing the execution of micro-hole craniotomy along a predefined trajectory. Through systematic *in vitro* testing, we show that the smart-control system can precisely detect craniotomy termination with a response time of 34.0 ± 16.4 ms and a distance error of 16.2 ± 17.2 μm (Fig. 3c), which could be further improved by upgrading the hardware system and the control method in the future.

**Figure 3.**
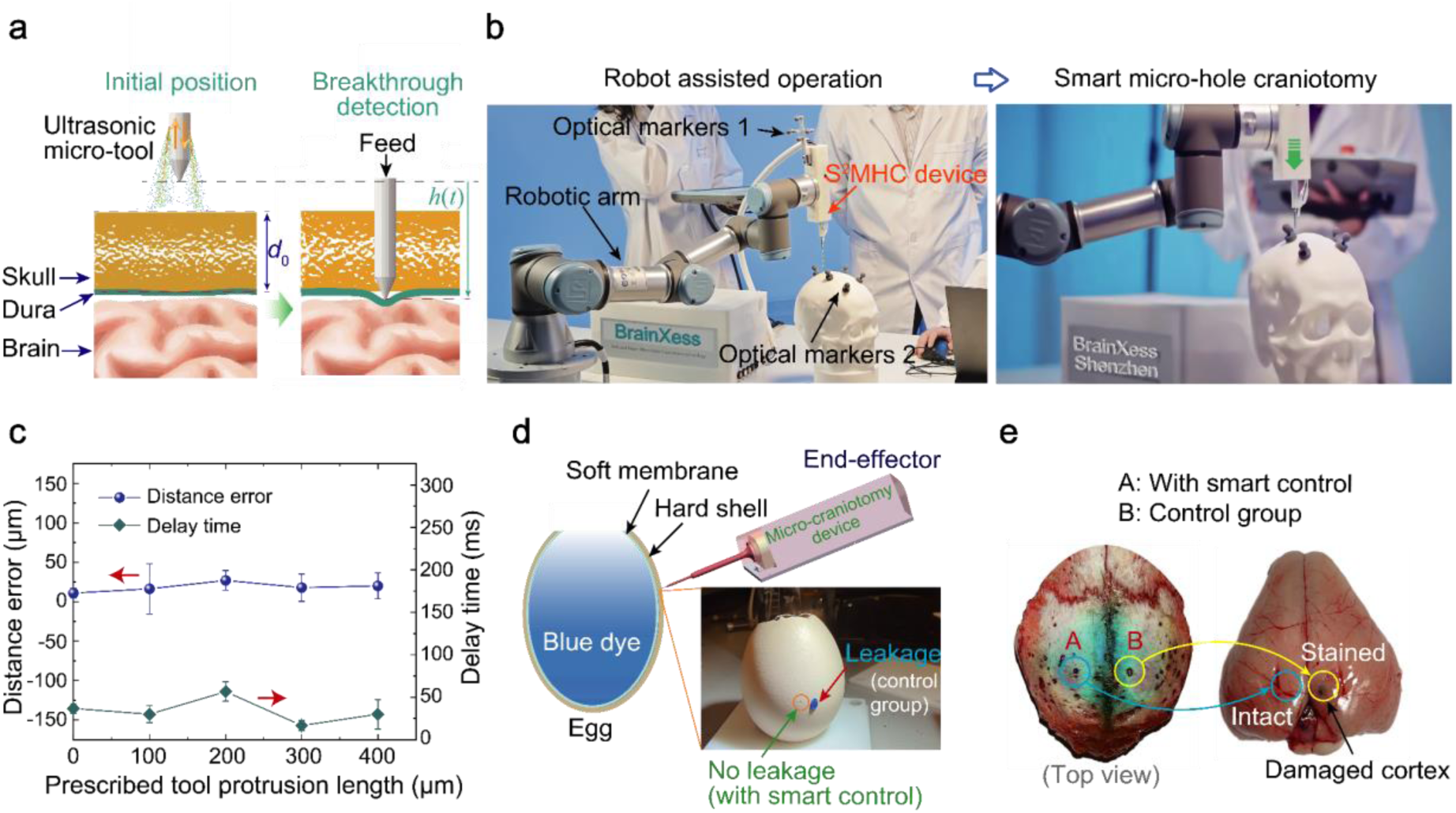
Smart control for safe micro-hole craniotomy at the skull-dura interface. (a) Detection of the craniotomy termination at the skull-dura interface. (b) Robot assisted micro-hole craniotomy; operated by a robot, micro-hole craniotomy can be performed freely at any location on the head along a pre-defined trajectory. (c) Performance of termination detection as represented by the average distance error and the delay time (*n* = 3), which were characterized by commanding the micro-tool to continue feeding for a prescribed distance after detection of craniotomy termination by the control system. (d) Demonstration of smart termination detection on a fresh egg. The fresh eggshell was filled blue dye solution. With smart control, a micro-hole was safely generated on eggshell without piercing the inner soft membrane. In the control group (no smart control), the soft membrane was damaged, leading to dye leakage. (e) Safe *in vivo* micro-hole craniotomy on New Zealand rabbits. After micro-hole craniotomy, blue dye was dispensed to the micro-hole; no staining was observed on the brain cortex with smart control.

Safe and precise micro-hole craniotomy was demonstrated by performing micro-hole perforation on a fresh eggshell (Fig. 3d). With smart control, the ultrasonic micro-tool was able to precisely penetrate the hard egg shell without piercing the underlying soft membrane. Safe micro-hole craniotomy with smart termination detection was further validated *in vivo* on anaesthetized rabbits. After micro-hole craniotomy, blue dye was dispensed into the micro-hole. In the control group, a blue staining was observed on brain, reflecting that continuous tool feeding without smart control would damage the dura and the cortex. In the test with smart termination detection, no blue staining was observed on the brain, indicating that the dura mater was intact (Fig. 3e).

### The minimally invasive hybrid BCI based on micro-hole craniotomy

Fig. 4a illustrates the concept of a minimally invasive hybrid BCI. Using the micro-hole craniotomy method, micro-holes can be safely prepared at specific locations on the skull. Micro-electrodes can then be implanted into these micro-holes, with the distal end contacting the dura and the proximal end placed subcutaneously at the scalp-skull interface. Once the acupuncture-like micro-wounds on the scalp have healed, a minimally invasive and hybrid BCI can be established by placing a wearable headset over the microelectrode implantation site (Fig. 4a). The microelectrode implanted in the skull establishes a stable and low-resistance pathway for measuring the neural signals in the brain or delivering electrical stimulation into the brain, providing an effective measure to overcome the constraints imposed by the skull.

**Figure 4.**
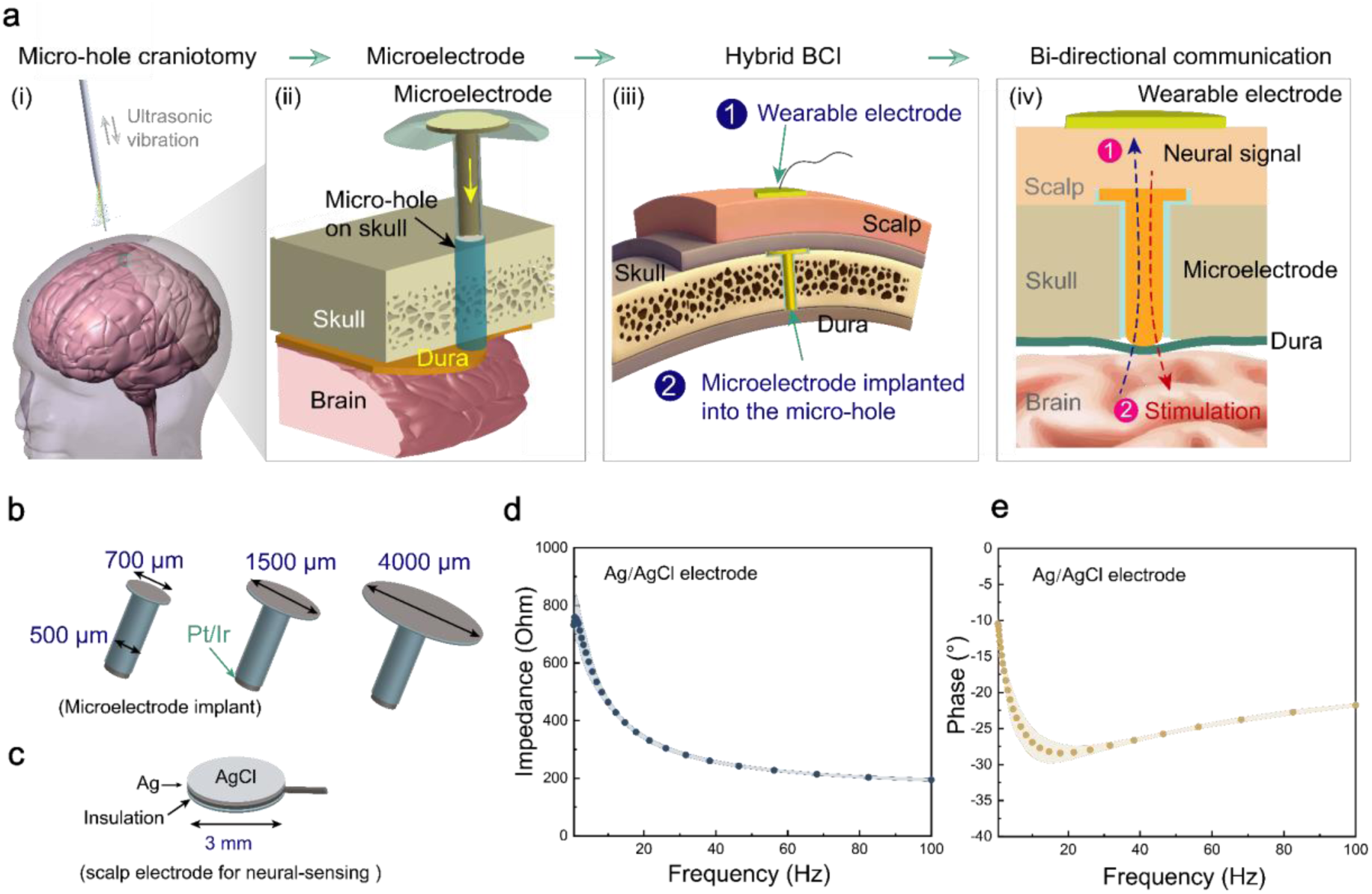
The minimally invasive hybrid BCI based on micro-intervention. (a) Implementation of the minimally invasive BCI, including micro-hole craniotomy, implantation of microelectrodes into micro-holes in the skull, and placement of wearable electrodes/headset on the scalp to establish the hybrid BCI. (b) Microelectrodes for implantation into micro-holes in the skull; the proximal end comes in sizes ranging from 700 to 4000 μm; the electrode shank has a diameter of 500 μm. (c) Customized Ag/AgCl electrodes for neural sensing on the scalp. (d-e) Impedance and phase angle spectrum of the Ag/AgCl electrode.

To verify the benefits and feasibility of the minimally invasive hybrid BCI, we prepared customized microelectrodes for micro-hole implantation. As shown in Fig. 4b, the Pt-Ir microelectrode has a cylindrical body with a diameter of 500 μm (the same as the micro-hole) and a flange with a diameter of 700-4000 μm at its proximal end as the depth-limiting structure. For animal experiments, a small Ag/AgCl electrode with a diameter of 3.0 mm was customized to acquire neural signals from the scalp of SD rats (Fig. 4c). This electrode was shown to have a low impedance (< 1 kΩ) in the 0-100 Hz frequency range and significant non-polarization characteristics, with the phase angle stabilizing between −10° and −30° in the weak capacitive range (see Fig. 4d-e). These characteristics (low impedance and high charge transfer efficiency) effectively reduce the thermal noise and the baseline drift introduced by the electrode-skin interface, ensuring a high signal-to-noise ratio and a stable baseline during neural signal acquisition.

### High-quality neural recording by the hybrid BCI

Effectiveness of the hybrid BCI for high-quality neural sensing was verified by measuring the resting-state neural signals and the somatosensory evoked potentials in response to a mechanical puncture. In the in vivo tests, microelectrodes (Fig.4b) were subcutaneously implanted into micro-holes generated in the rat skull (as E1 and E2, as indicated in Fig. 5a). E1 and E2 were located 6 mm apart anteriorly and posteriorly and positioned 3 mm from the midline, in rat’s somatosensory and visual cortical regions, respectively. C1 and C2, which were located symmetrically on the opposite side of the skull to the electrode implantation sites, were used as the control for acquiring EEG signals. Once the scalp wound had healed, the hairs were removed and the Ag/AgCl electrode (see Fig. 4c) were attached to the scalp at E1, E2, C1 and C2 with the conductive paste in order to non-invasively measure the neural signal. The hybrid BCI with implanted electrodes of proximal sizes of 0.7 mm, 1.5 mm, and 4.0 mm is denoted as the hybrid BCI-0.7, hybrid BCI-1.5, and hybrid BCI-4.0, respectively.

**Figure 5.**
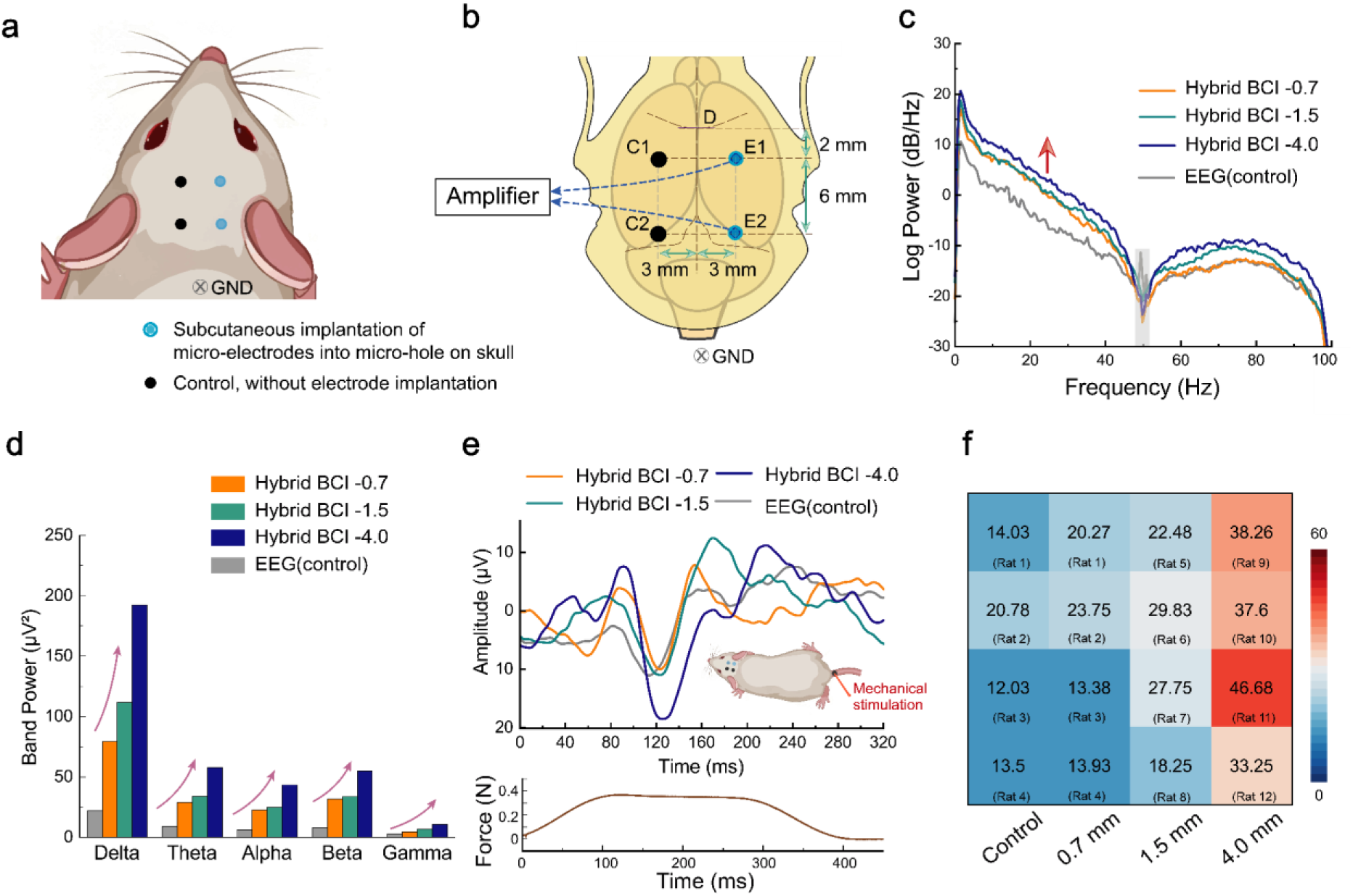
The hybrid BCI for high-quality neural recording. (a-b) Illustration of microelectrode implantation in the skull of SD rats. E1 and E2 are the locations for subcutaneous implantation of microelectrodes, as shown in Fig. 4b; C1 and C2 are the positions for EEG recording without electrode implantation (control). Neural signals were obtained by measuring the difference between E1 and E2 or C1 and C2, with reference to the common ground (GND). (c) Power spectral density (PSD) plots of the resting-state neural signals measured on the scalp of anesthetized rats (E1-E2) using the hybrid BCI with microelectrodes of different proximal dimensions (0.7 mm, 1.5 mm and 4.0 mm); control: the conventional resting-state EEG signals measured at C1-C2 without implanted microelectrodes. (d) Comparison of the power density in different frequency ranges for neural signals measured by different strategies as shown in (c). (e) Comparison of the somatosensory evoked potential waveforms measured by different methods. Mechanical puncture was applied to the rat tail using a blunt-tipped needle with a diameter of 0.3 mm that was controlled by a numerically controlled linear actuator. A typical force curve applied during the mechanical stimulation is also provided. (f) Signal-to-noise ratio (SNR) of somatosensory evoked potentials measured by the methods shown in (e). Neural signals for the control group in c-f were obtained at the C1-C2 locations in rats with implanted microelectrodes of a proximal size of 0.7 mm.

The hybrid BCI can evidently increase the strength of the measured neural signals in both the low- and high-frequency ranges. Fig. 5c shows the power spectral density plots of the neural signal measured using the hybrid BCI with implanted microelectrodes of different proximal dimensions (control: the conventional EEG measured between C1 and C2). Even the hybrid BCI using an implanted micro-electrode with a proximal size of 0.7 mm (hybrid BCI-0.7) could evidently increase the power of the measured neural signal in the low-frequency range (0-50 Hz). The hybrid BCI-1.5 increased the power further in the high-frequency range (50-90 Hz), and the hybrid BCI-4.0 increased the power across the entire frequency range (Fig. 5c). Furthermore, the neural signals were characterized into five typical frequency bands: delta (1-4.0 Hz), theta (4-8 Hz), alpha (8-13 Hz), beta (13-30 Hz), and gamma (30-100 Hz). As shown in Fig. 5d, the absolute power for the hybrid BCI groups was significantly higher than that for the control group. Compared to the EEG signals at different frequencies, the power strength of neural signals measured by the hybrid BCI-1.5 was 5, 3.8, 4.1, 4.1 and 2.6 times higher in the frequency bands of delta, theta, alpha, beta, and gamma, respectively. The power strength of the neural signals measured by the hybrid BCI-4.0 was 8.9, 6.3, 6.6, 6.7 and 4.1 times greater than that of the conventional EEG in the frequency bands of delta, theta, alpha, beta, and gamma, respectively.

The hybrid BCI enables the detection of evoked neural responses with a high signal-to-noise ratio. To demonstrate this, periodic mechanical stimulation was applied to the rat tail using a blunt-tipped needle to evoke the somatosensory evoked potentials (SEP), which were simultaneously measured between E1 and E2 using the hybrid BCI and between C1 and C2 using the conventional EEG. Fig. 5e shows the SEP waveforms measured by different methods after time-domain superposition and averaging. Compared to the control group (SEP measured by EEG), the SEP signals measured by the hybrid BCI group exhibited characteristic peaks with higher amplitudes. Therefore, the use of implanted microelectrodes with a larger proximal size can help increase the sensitivity in capturing functionally synchronized burst signals.

The hybrid BCI has been shown to function consistently well in repeated tests. The SEP tests were performed on 12 independent SD rats (four in each group) using the hybrid BCI. The signal-to-noise ratio (SNR) of the SEP measured on the different rats is compared in Fig. 5f. The data matrix exhibited a clear and consistent increase from the left to the right, from the conventional EEG to the hybrid BCI. In the EEG control group, the SNR for SEP ranged from 12.03 to 20.78; for the hybrid BCI-1.5, it increased to 18.25-29.75; and for the hybrid BCI-4.0, it increased further to 33.25-46.68. Therefore, despite the physiological differences between different animals, the hybrid BCI can reliably increase the strength of the neural signals measured (Fig. 5c-d) and increase the SNR for detecting evoked action potentials (Fig. 5e-f).

### Steady-state visual evoked potentials detected by the hybrid BCI

The hybrid BCI enhances the ability to reliably measure steady-state visual evoked potentials (SSVEP) with a high signal-to-noise ratio. Fig. 6a illustrates the experimental setup for evaluating the performance of the hybrid BCI in capturing SSVEP elicited by a light stimulator flashing at different frequencies. In the setup, an anesthetized rat was placed in a fully dark and electromagnetically shielded environment, with a white-light source positioned in front of the rat and flashing at different frequencies. As shown in Fig. 6b, the SSVEP was measured simultaneously between E2-Ref using the hybrid BCI method and between C2-Ref using the conventional EEG (the Ref electrode, located on the midline of the occipital bone, was used as the differential reference). Neural signals measured by different methods in response to the periodic flashing light stimuli (8 Hz, 10 Hz, 12 Hz, and 14 Hz) were processed to extract the frequency-domain features.

**Figure 6.**
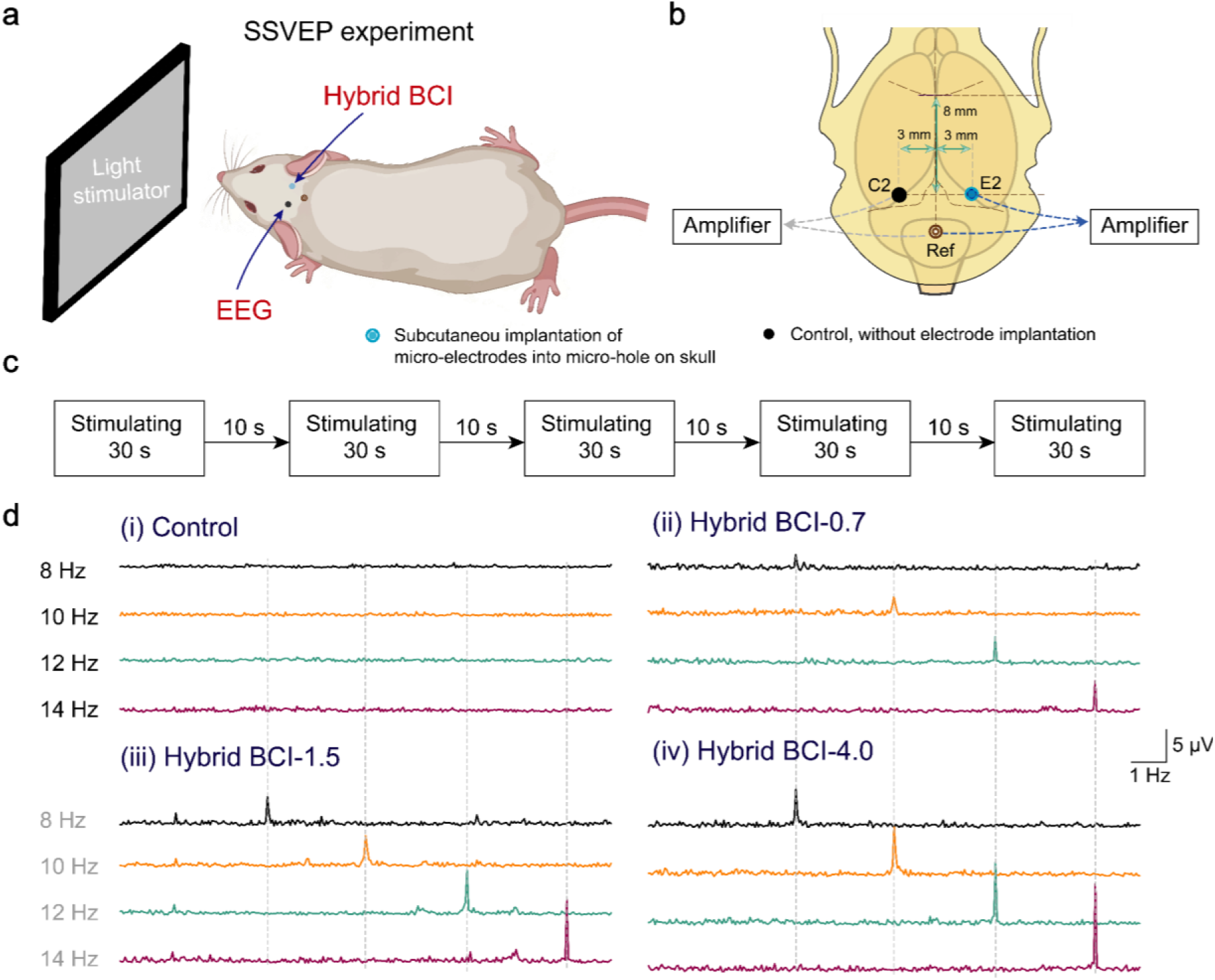
Tests on steady-state visual evoked potentials (SSVEP) measured by different methods. (a) Schematic diagram of showing the SSVEP experimental setup. (b) SSVEP was measured between E2 and Ref for the hybrid BCI group or between C2 and Ref for the control group. (c) The protocol for the SSVEP test; the SSVEP was elicited by the light stimulator flashing at different frequencies with a constant pulse width of 900 μs. (d) Comparison of the SSVEP measured by different methods. Neural signals for the control group were obtained at C2 for rats implanted with the microelectrode with a proximal size of 0.7 mm.

Figure 6d shows a comparison of the power spectrum plots of the SSVEP elicited by flashing lights of different frequencies and measured by different methods. In the control group (the conventional EEG), the spectral plots of the EEG signal were relatively flat within the stimulation frequency range, with no evident characteristic energy peaks around the four testing frequencies (Fig. 6d(i)). This would be attributed to skull’s attenuation and filtering effects. This phenomenon highlights the inadequacy of traditional non-invasive scalp electrodes in capturing low-intensity neural activity. However, distinct SSVEP peaks could be observed at 8 Hz, 10 Hz, 12 Hz and 14 Hz in the hybrid BCI group (Fig.6d(ii-iv)). Even in the hybrid BCI-0.7 group, the implanted microelectrode with a proximal size of just 0.7 mm was able to capture distinct narrow-band peaks around the stimulation frequency. For the hybrid BCI-1.5 group, steep and prominent characteristic SSVEP peaks appeared at the stimulation frequencies of 8 Hz, 10 Hz, 12 Hz, and 14 Hz (or their resonant frequencies, not shown here), with a significant increase in the amplitude. As the proximal end of the implanted microelectrode increased to 4.0 mm, the intensity of the detected SSVEP increased further (Fig. 6d(iv)), providing high-quality signals for neural decoding or control purposes.

Stability of the hybrid BCI for SSVEP detection was evaluated via repeated tests on different animals (four animals per group). The merits of the hybrid BCI was quantitatively evaluated by comparing SNR of SSVEP on different animals under different stimulation frequencies in Fig. 7a. Meanwhile, Fig. 7b-e shows the frequency spectrum for neural signals measured by the different methods on four independent rats at stimulation frequencies of 8 Hz, 10 Hz, 12 Hz and 14 Hz using the fast Fourier transform method (the 30-second stimulation was repeated five times on each rat). In the control group (Fig. 7b), the SSVEP signals were difficult to detect across different animals, owing to the attenuation effects caused by the skull.

**Figure 7.**
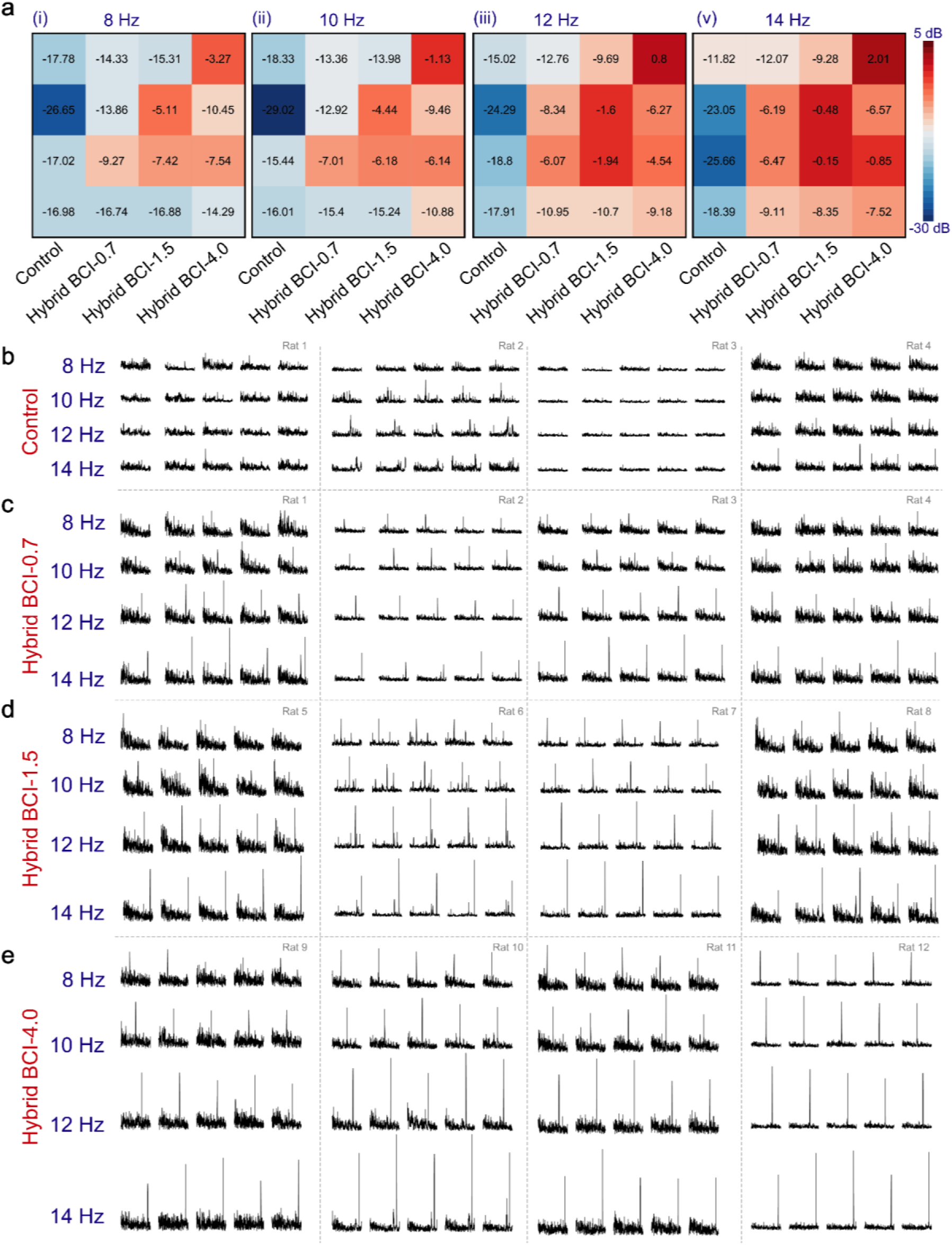
Repeated SSVEP tests performed using the hybrid BCI method. (a) Heatmap showing the SNR of the SSVEP measured using different methods under different conditions. (b-e) SSVEP spectra measured using the setup shown in Fig. 6a by the conventional EEG (control), the hybrid BCI-0.7, the hybrid BCI-1.5, and the hybrid BCI-4.0. Neural signals for the control group were obtained at C2 from rats implanted with a microelectrode of a proximal size of 0.7 mm. The SNR value in (a) is an average of the five data points shown in b-e for each animal. The tests were performed on four different animals under each condition. In (a), each block represents an independent rat as in Fig. 5f.

When the 0.7 mm microelectrode was implanted, the SSVEP could be clearly detected across four different animals, despite the fluctuations in SNR (see Fig. 7c). In the hybrid BCI-1.5 and the hybrid BCI-4.0 group, pronounced SSVEP peaks with a higher amplitude were detected with excellent inter-subject consistency. In particular, the hybrid BCI-4.0 group exhibited consistent observation of high-intensity SSVEP characterized by sharp, narrow-band peaks. Additionally, the SNR of the SSVEP measured by the conventional EEG remained low, with no significant difference observed across tests at different stimulation frequencies. Regarding the hybrid BCI, the SSVEP measured by the hybrid BCI tends to have higher peak amplitudes and SNR as the stimulation frequency increases. These findings further confirm the benefits of the hybrid BCI for acquiring high-quality neural signals for future brain-control applications based on this breakthrough strategy.

### Precise deep brain neural modulation enabled by the hybrid BCI

The skull poses a significant barrier for precise and localized neural modulation, particularly when the target is located deep within the brain. In the conventional transcranial electrical stimulation, due to skull’s high resistance, applied current would spread over a large area on the scalp before entering the brain via the path of least resistance. Only a very small portion (∼10%) of the applied current actually entered the brain^17^. It is also difficult to achieve precise brain stimulation via the conventional transcranial electrical stimulation. To address this issue, researchers have proposed non-invasive temporal interference stimulation (TI), which involves applying a pair of high-frequency currents applied to the scalp to generate a low-frequency envelope electric field within the brain. This enables electrical stimulation of deep brain targets^36,37^.

Due to the spreading effect of the electrical current and the skull’s inherent heterogeneous properties, there are two issues that need to be overcome for clinical application of TI, insufficient stimulation strength and low stimulation accuracy. Increasing the current applied to the scalp to strength the stimulation intensity would cause safety issues such as skin burns. Due to the wearable nature of TI stimulation, it is almost impossible to ensure that the electrodes are applied exactly at the same location on the scalp between two applications. Any shift in the location of wearable electrodes will exaggerate the targeting error of the enveloped electric field in the deep brain. Therefore, despite the safety and ease of use of wearable BCI, the shielding and attenuation effects of the skull present an insurmountable physical barrier to precise electrical stimulation of the brain.

To highlight the benefits of the hybrid BCI for precise neural modulation, the capability of the hybrid BCI for precise whole-brain stimulation was investigated via simulation and compared with the conventional transcranial temporal interference stimulation (TI). The simulation was performed on a simplified spherical model with a size of the normal human head^38,39^, comprising the scalp, the skull, the cerebrospinal fluid and the brain (Fig. 8a). Two typical scenarios were compared in the simulation, including the conventional transcranial stimulation (either the transcranial alternating current stimulation (tACS), or the transcranial temporal interference stimulation (tTI), Fig. 8b).

**Figure 8.**
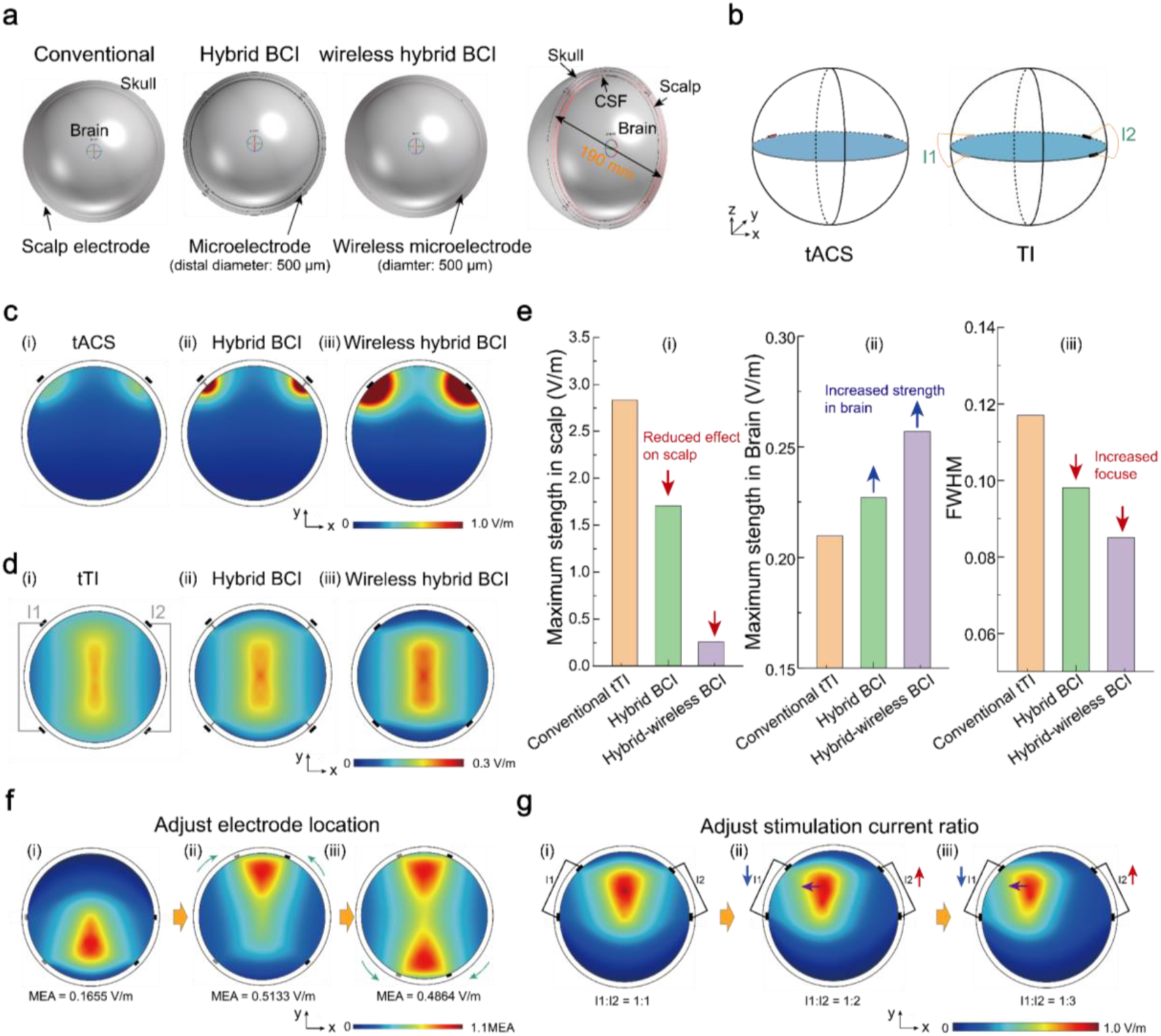
The potential of the hybrid BCI for precise stimulation of deep brain targets across the whole brain. (a) The simulation model on a simplified brain model, showing the electrode configuration for different scenarios: the conventional transcranial electrical stimulation, the hybrid BCI with implanted microelectrodes and wearable electrodes attached on the scalp, and the hybrid wireless BCI with implanted wireless microelectrodes. (b) Illustration of the transcranial alternating current stimulation (tACS) with two electrodes, and the temporal interference stimulation (TI) with four electrodes. (c) Simulation of the electrical field distribution in the brain for alternating current stimulation by different strategies. (d) Simulation of the interferential electric field envelope modulation in the brain by different methods. (e) Comparison of the maximum electrical field strength induced in the scalp or brain by different methods; the full width at half maximum (FWHM), which is the distance between two points at half the peak intensity in the *x* direction in Fig. 8d, represents the focus of the envelope electric field. (f) The focus of the envelope electrical field can be adjusted by changing the electrode locations; the values are normalized to the maximum envelope amplitude (MEA); color maps are in V/m. (g) The focus of the envelope electrical field for neural modulation can be adjusted by changing the current ratio between the two pairs of electrodes. I1 = I2 = 1 mA in c, d and f; I1+I2= 2 mA in g; the color maps in c,d, f and g are in V/m.

The ultimate version of the hybrid BCI, the wireless hybrid BCI, is also included for comparison purposes. The wireless hybrid BCI will have wireless microelectrodes implanted in the skull, which are wirelessly powered by and coupled with a wearable headset to enable wireless neural recording or multi-mode neural modulation. With the wireless hybrid BCI, electrical stimulation can be directly applied to the dura, bypassing the skull and the scalp to achieve effective and precise neural modulation.

Fig. 8c illustrates the electrical field distribution in the brain resulting from the conventional tACS, the hybrid BCI (with implanted microelectrodes and scalp electrodes), and the wireless hybrid BCI (with implanted wireless microelectrodes and a wearable wireless headset). In the conventional tACS, the skull’s attenuation effects cause a significant amount of electrical current to be shunted laterally within the scalp layer. This results in an effective electric field reaching the brain that is quite weak (Fig. 8c-i). In comparison, in the hybrid BCI group, implantation of microelectrodes into the skull creates low-resistance pathways for current flow and greatly minimizes the shielding effect of the skull, significantly increasing the stimulation strength in the brain (the cortex) is significantly increased (Fig. 8c-ii). For the wireless hybrid BCI, when the stimulation current is applied on the dura (with wireless power transmission and wireless control), electrical current’s spreading effect within the scalp can be eliminated, increasing the maximum stimulation strength (Fig. 8c-iii).

We demonstrate the feasibility of precise deep brain modulation using the hybrid BCI. Fig. 8d shows a comparison of the interferential electric field envelope amplitude maps of the conventional transcranial TI (tTI), the hybrid BCI, and the wireless hybrid BCI. Fig. 8e shows a comparison of the maximum envelope amplitude (MEA) in the scalp or the brain. The focus of the envelope electric field is represented by the full width at half maximum (FWHM), which is the distance between two points at half the peak intensity in the x direction (see Fig. 8d). A smaller FWHM indicates a more focused envelope field. Similar to the tACS, the transcranial TI has a low envelope amplitude in the brain, with a diffused center region (Fig. 8d-i). For the TI stimulation using the hybrid BCI, the induced envelope electrical field is more focused, with a larger MEA in the center. When the TI stimulation is performed using the wireless hybrid BCI, neural modulation of a higher intensity can be applied precisely to a focused zone with an even smaller FWHM. Findings in Fig. 8d-e clearly demonstrate the advantages of the hybrid BCI over the conventional non-invasive TI for effective and selective stimulation of deep brain targets.

The potential of the hybrid BCI for precise neural stimulation across the whole brain is further demonstrated. In Fig. 8f-g, we show that the focus of the envelope electric field can be flexibly controlled by adjusting either the electrode implantation locations (as in Fig. 8f) or the current ratio between I1 and I2 (as in Fig. 8g). Furthermore, the microelectrodes can be implanted at any predefined locations across the head and optimized by an AI-agent for the minimally invasive BCI. In this manner, one or more targets located deep in the brain can be stimulated in a specific spatial and temporal order, providing a precise, robust and versatile and tool for brain network modulation.

## 3. Conclusion and discussion

In this study, we introduce a minimally invasive hybrid BCI architecture comprising of four elements: the safe and smart micro-hole craniotomy; distributed microelectrodes implanted in micro-holes in the skull; a bi-directional wearable headpiece; and an AI-assisted planning and control framework. The distributed microelectrodes are implanted subcutaneously at pre-defined locations in the skull, with their distal ends in contact with the dura and their proximal ends at the scalp-skull interface. The wearable headpiece is coupled with the implanted microelectrodes for neural recording or stimulation. The AI-assisted brain agent is intended to support the deployment of the hybrid BCI across the whole brain, including but not limited to the planning of the electrode implantation, optimizing the pairing between implanted microelectrodes and the wearable headset, or decoding neural signals for closed-loop control. The hybrid BCI thus establish a technological pathway between the conventional non-invasive and the intracranial invasive BCI systems.

The enabling technique of the hybrid BCI is the ultrasonic vibration assisted micro-hole craniotomy method. Currently, achieving safe micro-hole craniotomy without impairing the dura is difficult using existing techniques. In this study, the safe and smart micro-hole craniotomy method developed by our team enables micro-hole generation in a convenient manner like acupuncture. Micro-tunnels can be dexterously prepared at pre-defined positions on the skull with a low axial force, minimal thermal effects, and precise termination detection. This significantly reduces the risk of surgical inflammation and provides a micro-tunnel to reach the skull-dura boundary while maintaining dura integrity.

Feasibility of the hybrid BCI has been verified in vivo with rat experiments. We demonstrated that, in comparison with the conventional scalp EEG, microelectrodes placed in micro-holes in the skull could increase resting-state spectral power intensity and improve the signal-to-noise ratio for detecting the somatosensory and steady-state visual evoked responses. These in vivo results provide a solid proof that the hybrid BCI architecture can substantially enhance the access to brain-generated electrical signals across multiple frequency bands and response paradigms. The robust performance of the hybrid BCI has also been verified through repeated tests. The next step is to develop a wireless hybrid BCI and test it on free-moving animals for wireless neural recording and stimulation. Harvesting high-quality, high-fidelity neural signals in a minimally invasive manner will provide valuable information for future neural decoding and closed-loop brain control applications.

The distributed architecture provides a scalable pathway for whole-brain coverage. Distributed microelectrodes (or nodes) can be placed at selected locations on the skull, with more nodes incrementally added as required for clinical purposes. Whole-brain coverage is difficult to achieve by existing invasive BCI methods. While it is technically possible to implant as many electrodes as needed, such an invasive procedure may not be clinically feasible. The hybrid BCI therefore offers a versatile means of interacting with the brain via distributed microelectrodes implanted in micro-holes across the cranial surface, the number and position of which can be dynamically adjusted according to patient’s needs.

Our simulation results show that the skull-dura microelectrodes can reshape the current pathways, reduce superficial current shunting, and improve both the electric-field strength and the focality for stimulating deep brain targets, compared to the conventional transcranial stimulation. A simplified spherical model was used to enable a controlled comparison of different stimulation strategies. Together with the programmable electrode locations and current ratios, the simulation results confirm the feasibility of steering stimulation towards cortical and deep targets across the brain, providing a solid basis for future multi-target, temporally coordinated network modulation. Thus, the hybrid BCI will provide an effective and well-controlled tool for whole-brain modulation, which is highly demanded for the treatment of neurodegenerative and neuropsychiatric diseases, including but not limited to refractory depression, addiction, neuropathic headaches, sleep disorders and Alzheimer’s disease. Extending this hybrid framework to include patient-specific head models, realistic electrode-tissue interfaces, dose-response study and in vivo validation will define the translational performance of the hybrid BCI in the near future.

In the next stage, these validated building blocks will be integrated into a chronic wireless hybrid BCI system with bidirectional reading and writing functionalities, while preserving the core architecture of a simple implanted interface coupled to an externally upgradable wearable platform. Priority studies will include investigating long-term electrode-dura stability, biocompatibility, the safety of repeated-stimulation, MRI compatibility, artifact recovery during read-write switching, and closed-loop performance. These are tractable engineering and translational milestones that can be achieved through large-animal studies, verification testing, and clinical feasibility trials.

From a clinical application perspective, the microelectrodes of the hybrid BCI can be implanted in a manner similar to hair transplant, which makes them readily accepted by a large patient population. The microelectrodes can be designed to be MRI-compatible for long-term implantation in the skull. The hybrid nature of this system allows it to evolve at both the hardware and software levels, either by implanting more electrodes or by upgrading the wearable headset and the AI agent.

Taken together, the present study validates the foundational architecture of the minimally invasive hybrid BCI at three complementary levels: safe and controllable access to the brain via micro-holes in the skull, enhanced neural sensing in vivo, and computationally supported modulation of deep brain targets. The convergence of surgical, electrophysiological, and computational evidence establishes a clear and credible translational pathway towards a chronic, wireless, distributed, and bidirectional BCI system. Through ongoing validation, this hybrid BCI architecture will occupy a distinct design space between the conventional noninvasive BCI and existing invasive BCI in which the burden of the surgical procedure, the signal quality, the spatial extensibility, and the system’s long-term upgradability are jointly optimized, to support high-quality cortical recording, precise deep brain modulation, and network-level closed-loop control.

## 4. Methods

### Simulation of the micro-hole craniotomy

The micro-hole craniotomy process was simulated in Abaqus using the smoothed particle hydrodynamics (SPH) method^30^. A freshly harvested skull bone sample from the adult New Zealand rabbit (12-14 weeks) was scanned with the Micro-CT (SkyScan 1176, Bruker, resolution: 8.8 μm) ^40^. A representative CT image was used to construct skull bone’s 2D model. In order to mimic skull bone’s micro-porous structure, micropores were randomly generated in the selected CT image using Matlab^58,59^; the cortical bone was set to have a porosity of 7.8%^41^; the resulted image was subsequently processed by Mimics and SolidWorks, generating a two-dimensional executable file which was subsequently processed in Abaqus for particle segmentation with a customized Python script^42^. A 2D SPH model was then constructed. The ultrasonic micro-tool was modelled as a rigid body; side surfaces of bone sample were fully constrained; skull sample’s bottom surface was constrained in all directions except for the area around the hole exit. Skull tissue was assumed to be isotropic^43^. Elastic and plastic properties of the skull bone were adopted from published studies^44–46^; the shear damage model was used to simulate skull bone’s damage behavior. Stress, plastic strain, material flow around the micro-tool were simulated.

### Micro-hole craniotomy experiment

Micro-hole craniotomy experiment and finite indentation tests were performed. Animal experiments were performed with institutional approval (IACUC approval number: SYSU-IACUC-2023-000446). Bone samples from central part of the skull (length: 12 mm; width: 6 mm; thickness: ∼ 1.5 mm) were harvested from euthanized adult New Zealand rabbits (18-21 weeks) and preserved in PBS solution prior to experiments. Micro-hole craniotomy was performed by feeding the ultrasonic tool towards the skull bone; the craniotomy force was recorded by a force sensor. After experiment, bone samples were freeze-dried, sputtered-coated and observed under SEM. To investigate the micro-hole craniotomy process, the micro-tool was fed to a finite depth in skull (100-1000 μm) and then retracted to observe the hole morphology under SEM.

### Low force micro-hole craniotomy in comparison with skin penetration

In order to highlight the merit of low-force micro-hole craniotomy, *in vivo* and *in vitro* skin penetration tests were performed with a subcutaneous injection needle (bevel-tipped; diameter: 500 μm). *In vivo* tests were conducted on anesthetized SD rats (12-14 weeks). After hair removal, skin penetration was executed on rat back with numerical control. *In vitro* tests were performed on the skin of anesthetized rats. The maximum skin penetration force was recorded for comparison. Animal experiments were performed with institutional approval (SYSU-IACUC-2023-000446).

### *In vitro* tool-dura contact test

Dura samples were harvested from euthanized adult New Zealand rabbits and clamped onto a customized sample holder. The ultrasonic tool was lowered to press the dura mater for a distance of 500 μm, mimicking tool-dura contact at the end of craniotomy; after different time of tool-dura contact (2-60 seconds), tool-dura contact area was observed under the optical microscope and characterized under SEM (after freeze-drying and sputter coating with gold).

### Micro-hole craniotomy with low thermal effects

As tool-bone friction would be the main cause for heat generation in the micro-hole craniotomy process, effective cooling was applied to reduce the temperature during micro-hole craniotomy. Saline coolant (temperature: 20℃) was applied to the craniotomy site. Temperature in skull bone around the micro-hole were systematically investigated under different cooling conditions with micro-thermal couples which were pre-embedded in pre-drilled micro-holes of the bone sample. Micro-hole craniotomy was performed at different distances off the micro-thermal couple.

Temperature profile around the micro-hole was simulated with the steady-state heat transfer model in Abaqus. A 500 μm diameter micro-hole was modelled in a rectangular skull bone sample (thermal conductivity: 0.56 W/(m·K)^47^). Consistent with the *in vitro* test, temperature for bone sample’s top surface was set to be 20°C; temperature for bone surfaces immersed in water or in air was 37°C and 25.4℃ (room temperature measured at the day of experiment), respectively. An initial temperature was applied to micro-hole’s inner surface to simulate the temperature profile; mean difference between simulated temperature and experimentally measured temperature at different distances off the hole wall (0.5, 1.0, 1.5, and 2.0 mm) was calculated for each initial temperature. Temperature profiles in bone tissue around the micro-hole with or without cooling were simulated and compared with experimental results.

### Fast recovery after micro-hole craniotomy via *in vivo* tests

Quick recovery after micro-hole craniotomy was investigated *in vivo* on adult SD rats (6-8 weeks). After proper anesthesia, hair removal and skin disinfection, a small incision was made on scalp for micro-hole craniotomy with optimized cooling (control: no cooling). Four micro-holes were symmetrically generated on rat skull (two on the left and two on the right; hole-to-hole distance: ∼ 4 mm); after experiment, the incision on scalp was properly sutured. At 2-5 weeks after surgery, rats were euthanized to harvest skull samples for H&E staining^48^. Animal experiments were performed with institutional approval (IACUC approval number: SYSU-IACUC-2023-000446).

### Smart control for craniotomy termination detection

A smart control system has been developed for precise detection of craniotomy termination. A control algorithm is developed for the detection of tip breakthrough or craniotomy termination; skull thickness *d*_0_ along the craniotomy path could be used as the reference. Performance of the craniotomy termination detection system was investigated from the perspective of delay time (or response time) and distance error. Systematic *in vitro* tests were performed with fresh rabbit skull samples. The delay time is defined as the time difference between the moment of breakthrough detected by the control algorithm to the moment of actual termination of tool feeding (recorded by the control algorithm). As for the distance error, the ultrasonic micro-tool was commanded to continue feeding for an extra distance of *L*_a0_ after breakthrough detection; micro-tool’s actual protrusion length over skull’s inner surface (*L*_a1_) minus the prescribed protrusion length (*L*_a0_) was defined as the distance error (*L*_a1_ and *L*_a0_ were measured under the microscopic camera).

*In vitro* demonstration of smart termination detection was performed on fresh eggs. An opening was made on the eggshell to fill the shell with blue dye solution. Micro-hole perforation was performed on eggshell using the ultrasonic micro-tool with the smart control method (control: without smart control). If shell breakthrough could be accurately detected, eggshell’s inner membrane would be intact, and dye leakage would not be observed. Micro-holes generated on eggshell were observed under the optical microscope and SEM to characterize the integrity of the soft membrane after micro-hole perforation.

Micro-hole craniotomy with smart control was demonstrated *in vivo* on New Zealand rabbits. Animal experiments were performed with institutional approval (IACUC approval number: SYSU-IACUC-2023-000446). After proper anesthesia, hair removal and skin disinfection, an incision was made on scalp to expose the skull; two micro-holes were symmetrically performed on skull (one on the left, and one on the right). In the experiment group, micro-hole craniotomy was performed with smart control and cooling. In the control group, micro-hole craniotomy was executed with disabled smart control, allowing the ultrasonic tool feed to a depth of 4.0 mm beneath skull surface. After micro-hole craniotomy, blue dye solution (Evans blue) was dispensed into micro-holes on the skull. 30 minutes later, the rabbit was euthanized; the skull was removed to examine (1) whether a through-hole was formed on skull and (2) integrity of the brain cortex beneath the craniotomy site (if the brain was damaged, impaired cortex would be stained by the blue dye). A through-hole and intact cortex would confirm the safe and smart micro-hole craniotomy.

### Microelectrode implantation

Animal experiments were performed with institutional approval (SYSU-IACUC-2026-000788). Twelve male SD rats (250-300 g) were randomly assigned into 3 groups *(n* = 4 per group). During surgery, animals were anesthetized with isoflurane (induction 3%-5%, maintenance 1%-3%) and secured onto a stereotaxic frame; meloxicam (1 mg/kg) was subcutaneously administered for preemptive analgesia. Micro-hole craniotomy was performed at specific locations shown in Fig. 5b or Fig. 6b. After hair removal, the surgical site was sterilized with iodopovidone and 70 v/v% ethanol. Microelectrodes with different proximal dimensions (0.7 mm, 1.5 mm, and 4.0 mm; see Fig. 4b) were implanted into the micro-holes (E1 and E2 in Fig. 5b), with electrode’s distal end in contact with the dura and its proximal end subcutaneously implanted at the skull-scalp interface. Length of the microelectrode was customized to fit the local skull thickness. Contralateral locations C1 and C2 were used for scalp EEG without micro-intervention^49^. After surgery, meloxicam (1 mg/kg) was subcutaneously administered daily for 3 consecutive days for postoperative analgesia, and all rats were monitored daily. Animals were allowed to recover under standard housing conditions for 1 week to allow the scalp incisions to heal. All twelve animals were properly recovered from the surgery for subsequent testing.

### Resting state neural signal

After anesthesia with isoflurane (1%-3%), hair removal and skin disinfection, the Ag/AgCl electrodes (Fig. 4c) were attached with conductive paste to C1 and C2, or E1 and E2; the common ground electrode (GND; Fig.5b) was placed on the skin at the posterior neck. Resting-state signals were recorded continuously for 5 minutes per session; signals were amplified using a four-channel differential amplifier (A-M 1700) and recorded as the differential potential between C1-C2 (control: scalp EEG) or E1-E2 (with microelectrodes implanted)^49,50^; the hardware band-pass filter (0.5–500 Hz) and the 50 Hz notch filter were used during data acquisition. Neural signals for the control group were obtained at C1-C2 only in rats implanted with the micro-electrodes with a proximal size of 0.7 mm. Neural signals were recorded via a data acquisition card (DAQ) at a sampling rate of 1000 Hz.

### Power spectral analysis

Power spectral density (PSD) was calculated using the Welch method with 2 seconds Hamming-windowed segments (overlapping ratio: 50%). PSD values were displayed in the logarithmic scale in the frequency range of 0-100 Hz. Band power was calculated by integrating the PSD within the delta (1-4 Hz), theta (4-8 Hz), alpha (8-13 Hz), beta (13-30 Hz), and gamma (30-100 Hz) ranges.

### Somatosensory evoked potentials (SEP)

Rats were anesthetized and placed in an electromagnetically shielded environment. Somatosensory stimulation was delivered to the based of the tail by advancing a blunt-tipped needle (diameter: 300 μm) for 2 mm using a numerically controlled, force-recording linear actuator. Prior to stimulation, the blunt-tipped needle was adjusted to be in contact with the skin. Stimuli were delivered periodically (one cycle of stimulation: 5 s); a total of 60 cycles of stimulation was applied per rat. A customized LabVIEW program synchronized actuator control, force acquisition, and neural signal recording.

Each stimulus was defined as t = 0 ms. A 500 ms neural signal epoch extending from −25 ms to 475 ms relative to the stimulus onset was extracted for each trial. The 60 epochs were time-aligned and averaged to suppress spontaneous background activity and to obtain stable SEP waveforms^51,52^. SEP signal quality was quantified using the signal-to-noise ratio (SNR) = 20 log_10_(*A*_signal_/*RMS*_noise_), where *A*_signal_ is the peak-to-peak SEP amplitude, and *RMS*_noise_ is the root-mean-square amplitude of the 25 ms pre-stimulus baseline^51^.

### The steady-state visual evoked potential (SSVEP)

Rats were anesthetized and placed in an electromagnetically shielded cage covered with blackout. The visual stimulation system consisted of a strobe controller and a white-light panel source (70 mm by 70 mm) that was placed in front of the rat eyes^53^. Onset, frequency, and pulse width of the light stimulation were precisely controlled by the strobe controller. In the test, four visual stimulation frequencies (8 Hz, 10 Hz, 12 Hz, and 14 Hz) were used. For each frequency, five 30 s stimulation blocks were delivered, separated by 10 s dark intervals.

The flash pulse width of the light stimulus was strictly controlled at 900 μs. For example, an 8 Hz condition delivered eight evenly spaced flashes per second. Frequency-domain responses were obtained using the fast Fourier transform (FFT) operation to quantify SSVEP components at stimulation frequencies^53–55^. SNR of the SSVEP was calculated as follows:

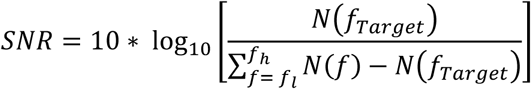

where, N(fTarget) denotes the power at the target frequency, and 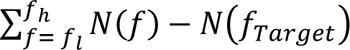 denotes the summed power within the 5-15 Hz band after exclusion of the target frequency component.

### Simulation of electrical stimulation

A simplified concentric spherical head model was constructed for simulation^38,39^ (Fig. 8a-b). The model was constructed in Solidworks and imported into COMSOL Multiphysics for analysis. The diameters for the scalp, skull, cerebrospinal (CSF) layer, and the brain were set to 190 mm, 182 mm, 172 mm and 170 mm, respectively. The wearable scalp electrode was modeled with a diameter of 2 mm and a thickness of 1 mm. The electrical properties for different tissue and the electrode were obtained from the literature^56–59^ (see Table 1); all materials were assumed to be homogeneous. Also, under the quasi-static approximation for kHz frequency electromagnetic fields, where the wavelength is significantly larger than the tissue dimensions, tissue properties were assumed to be frequency-independent^56^.

**Table 1.**
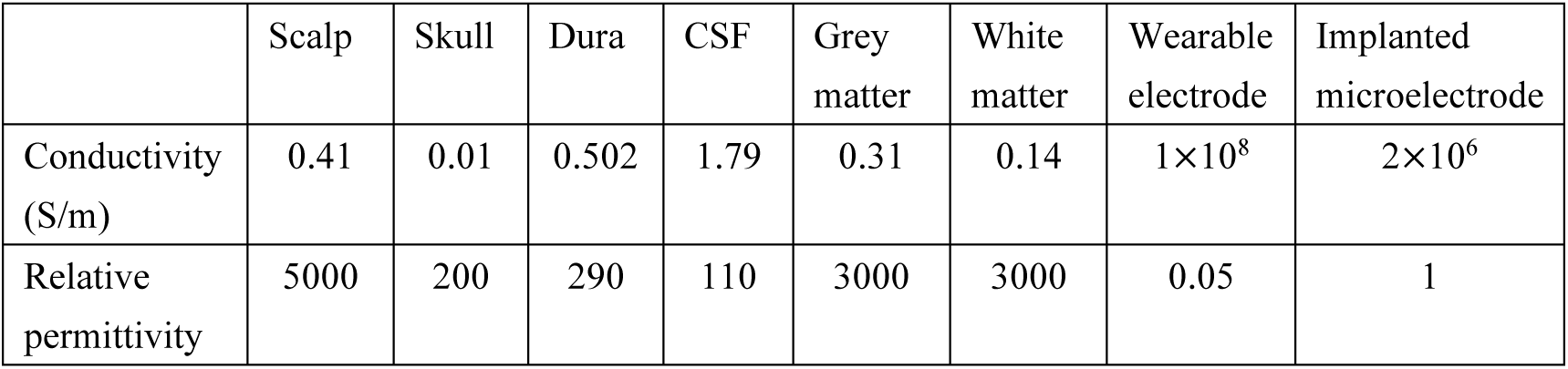
Material properties used in the simulation^56–59^.

|  | Scalp | Skull | Dura | CSF | Grey matter | White matter | Wearable electrode | Implanted microelectrode |
| --- | --- | --- | --- | --- | --- | --- | --- | --- |
| Conductivity (S/m) | 0.41 | 0.01 | 0.502 | 1.79 | 0.31 | 0.14 | $1 \times 10^8$ | $2 \times 10^6$ |
| Relative permittivity | 5000 | 200 | 290 | 110 | 3000 | 3000 | 0.05 | 1 |

The electrical field distribution in the brain generated by transcranial alternating current stimulation (tACS), temporal-interference stimulation (TI), the hybrid BCI, and the wireless BCI were simulated using the finite element method (FEM) in the electric currents interface of the AC/DC module in COMSOL Multiphysics following published studies^56,60^; electrical insulation boundary conditions were applied to the external surfaces of the model; Poisson’s equation, ∇·(*σ*∇*V*) = 0, was solved in the simulation, where *σ* is the electrical conductivity and *V* is the electrical potential. The electric field strength *E* was calculated as *E* = -∇*V*.

Regarding the electrode configuration, the wearable electrode was placed on the scalp in the conventional EEG. In the hybrid BCI, microelectrodes (diameter: 500 μm) were implanted in the skull with the distal end in contact with the dura and the proximal end at the scalp-skull interface. As for the wireless hybrid BCI, only the 500 μm diameter microelectrode was placed on the dura without the scalp electrode. In the simulation, electrodes were arranged on the *xy* plane (*z* = 0) at an angle of 45° to the center (Fig. 8b).

In the tACS or the hybrid BCI, the stimulation current was delivered through wearable scalp electrodes. In the wireless hybrid BCI, the stimulation current was delivered through the microelectrodes placed on the dura. In the simulation shown in Fig. 8c, d and f, I1 = I2 = 1 mA. In Fig. 8f, the electrode positions were adjusted to investigate their effect on the envelope strength. In Fig. 8g, the current ratio between I1 and I2 was adjusted while maintain a sum of I1+I2 as 2 mA.

The electric field vectors generated by a pair of electrodes were simulated. For the TI stimulation, the electric field vectors (*E*_1_ and *E*_2_) from the two electrode pairs (I1 and I2) were calculated separately. Envelope extraction for the TI was performed in MATLAB, and the maximum amplitude of the interference envelope 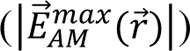 was computed analytically at each spatial position (*r*) using the following piecewise function:

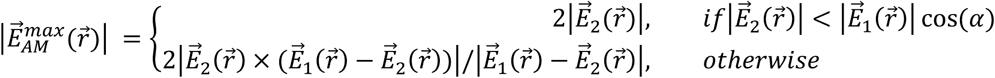

where, *α* is the angle between the two electric field vectors *E*_1_(*r*) and *E*_2_(*r*). The maximum envelope amplitude (MEA) was quantified in the brain tissue. Spatial focality was assessed using the full width at half maximum (FWHM), the distance between two points at half the peak intensity in the *x*-direction in the *xy*-plane (see Fig. 8d).

## Acknowledgements

Study on the minimally invasive BCI was supported by Shenzhen BrainXess Technology Co., Ltd. Correspondence and requests for materials should be addressed to Dr. Zhe Li at.

